# Transcriptomic landscape of *seedstick* in *Arabidopsis thaliana* funiculus after fertilisation

**DOI:** 10.1101/2023.11.13.566818

**Authors:** Maria João Ferreira, Jessy Silva, Hidenori Takeuchi, Takamasa Suzuki, Tetsuya Higashiyama, Sílvia Coimbra

## Abstract

In Angiosperms, the continuation of plant species is intricately dependent on the funiculus multifaceted role in nutrient transport, mechanical support, and dehiscence of seeds. SEEDSTICK (STK) is a MADS-box transcription factor involved in seed size and dehiscence, and one of the few genes identified as affecting funiculus growth. Given the importance of the funiculus to a correct seed development, allied with previous phenotypic observations of *stk* mutants, we performed a transcriptomic analysis of *stk* funiculi, using RNA-sequencing, to infer on the deregulated networks of genes. The generated dataset of differentially expressed genes was enriched with cell wall biogenesis, cell cycle, sugar metabolism and transport terms, all in accordance with *stk* phenotype. We selected eight differentially expressed genes involved with abscission, seed development or novel functions in *stk* funiculus, such as hormones/secondary metabolites transport, for transcriptome validation using qPCR and/or promoter reporter lines. Overall, the analysis performed in this study allowed delving into the STK-network established in Arabidopsis funiculus, fulfilling a literature gap. Simultaneously, our findings reinforced the reliability of the transcriptome, and identified processes and new candidate genes that will enable a better understanding on the role of this sporophytic structure and how seed development may be affected by it.

## Introduction

Seeds are intricate units essential to establish the next sporophytic generation of plants. In Angiosperms, the development of a seed starts with a key process called double fertilisation, where one sperm cell fuses with the egg cell, originating the diploid zygote, while the other fuses with the central cell, giving rise to the triploid endosperm (Hamamura *et al*., 2011). In the next stages of seed growth and maturation, an influx of nutrients, minerals, sugars and signals from the maternal plant through the vasculature of the funiculus is required (Becker *et al*., 2014). Finally, when the seed is mature, the formation of a seed abscission zone (SAZ) occurs, composed of a separation layer with thin cell walls that will degenerate at the end of the process and an adjacent layer with lignified cells (lignified layer), which produces the tension necessary for the separation of the tissues (Estornell *et al*., 2013). This process allows seeds to detach from the plant.

The funiculus is a specialised sporophytic structure that anchors the ovule to the placenta within the gynoecium of flowering plants, being the contact route for communication between them (Schneitz *et al*., 1995). It arises during ovule primordium and remains intact throughout ovule development until seed dehiscence occurs. Remarkably, there is a lack of information regarding *Arabidopsis thaliana* (*A. thaliana*) case-study compared to other plants such as *Brassica napus* (Tan *et al*., 2010; Chan and Belmonte, 2013; Chan *et al*., 2016) or *Phaseolus vulgaris* (Mawson *et al*., 1994), in which topics like anatomy or genetic regulation of funiculus development and underlying biological function have been extensively studied. So far, only one studied focused on clarifying these topics using *A. thaliana.* Khan *et al*. (2015) work showed using light microscopy that the funiculus is composed of an outer epidermis, a parenchymatous cortex, and an internal vascular core with xylem and phloem.

A comparative transcriptomic study between the zygotic regions and subregions of the seed and the funiculus in *A. thaliana* revealed the accumulation of several exclusive mRNAs in this stalk structure, presenting the funiculus as a transcriptionally distinct region from the other seed tissues (Khan *et al*., 2015). Of the 20 genes identified as funiculus-specific, some were associated with transport and others with carbon metabolism, but over 30% did not have an annotated function, thus, showing that regulatory networks in the funiculus require further investigation.

*SEEDSTICK* (*STK*) is one of the few genes identified to be involved in funicular patterning and growth (Pinyopich *et al*., 2003; Lan *et al*., 2023). It belongs to the MADS-box transcription factor (TF) family and is considered a master regulator of ovule development. Observations using a reporter line with the entire *STK* genomic region fused to a green fluorescent protein revealed high expression of this gene specifically in the ovules’ sporophytic tissues, in which the funiculus is included, from the beginning of ovule development until later stages when the mature seed is formed (Mizzotti *et al*., 2014). This expression pattern, together with many genetic-based studies performed over the years, supports the pivotal role of STK in orchestrating various processes during plant reproduction. STK redundantly controls ovule identity together with SHATTERPROOF1 (SHP1) and SHP2 (Pinyopich *et al*., 2003) and participates in transmitting tract development by interacting with NO TRANSMITTING TRACT (Herrera-Ubaldo *et al*., 2019) and CESTA (Di Marzo *et al*., 2020*b*). STK also determines fruit size by regulating cytokinin homeostasis and controlling *FRUITFULL* expression (Di Marzo *et al*., 2020*a*). Furthermore, STK is involved in seed development and size, affecting processes such as flavonoids biosynthesis (Mizzotti *et al*., 2014), cell wall organisation and remodelling properties (Ezquer *et al*., 2016; Paolo *et al*., 2021*a*; Di Marzo *et al*., 2022*b*,*a*) and cell cycle (Paolo *et al*., 2021*b*). Observations at floral stage 17 (Smyth *et al*., 1990) showed that *stk* mutants develop thicker and longer funiculi compared to wild-type (WT) and present defects in the SAZ, resulting in a decrease in the ability to dehiscence (Pinyopich *et al*., 2003). Balanzà *et al*. (2016) proposed a model with the STK network controlling the formation of the SAZ. According to this study, STK interacts with SEUSS co-repressor, forming a complex that will repress *HECATE 3*, enabling the correct deposition of lignin in the vasculature of funiculus cells. Nonetheless, only TFs have been reported as regulators for the establishment of the lignification pattern. However, in similar abscission processes occurring in the floral organ abscission zone (reviewed by Niederhuth *et al*., 2013) or silique dehiscence zone (reviewed by Yu *et al*., 2020), several intervenients besides TFs have been reported: hormones (like ethylene, auxin, jasmonic acid), small peptides and their receptors (INFLORESCENCE DEFICIENT IN ABSCISSION is perceived by HAESA or HAESA-LIKE 1), as well as cell wall modifying enzymes (cellulases, hemicellulases and pectinases).

Despite the significant achievements on STK role during seed shattering, many questions remain unanswered: Which downstream targets participate in seed abscission? Is STK controlling other processes in the funiculus? Is seed development being affected by them? Only one transcriptome on *stk* single mutant is currently available, however those data derive from inflorescences and fruits until five days after pollination, which do not include floral stage 17 (Smyth *et al*., 1990). Moreover, considering the different tissues contained in that transcriptome, it would be difficult to study, at the molecular level, processes happening in a specific tissue such as the funiculus.

In this study, we explore the transcriptomic data from *stk* and WT funiculi at floral stage 17, in order to investigate at the molecular level, the processes affected by *STK* absence in the funiculus. We provide information about enriched biological processes and molecular functions underpin the mutant funiculus phenotype, and we validate with qPCR and/or promoter reporter lines the expression of candidate genes, encoding proteins such as pectinases, laccases, or sugar transporters. Overall, we believe that by leveraging our results, novel players in the STK-mediated network in the funiculus can be found, and, importantly, whose functions may be affecting seed development and shattering.

## Material and Methods

### Plant material and growth conditions

Wild-type (WT) *Arabidopsis thaliana* (L.) Heynh. Columbia-0 seeds and the *stk-2* (Pinyopich *et al*., 2003) mutant line, referred to as *stk*, were sown directly on soil (COMPO SANA®, Germany) and kept for 48 h at 4 °C in the dark to induce stratification. Afterwards, all plants were grown under long-day conditions (16 h light at 22 °C and 8 h darkness at 18 °C) with 50-60% relative humidity and light intensity at 180 µmol m^-2^ s^-1^.

### Sample collection for RNA-sequencing

Under a SZ61 stereo microscope (Olympus), WT and *stk* siliques from stage 17 (st 17) flowers (Smyth *et al*., 1990) were first dissected, seeds were removed, and only the transmitting tract with the attached funiculi was placed into a 50 µL droplet of *in vitro* ovule culture medium (Gooh *et al*., 2015) in a plastic dish (Φ 3.5 cm). Each funiculus was separated using a surgical knife and transferred with a tungsten needle to another plastic dish containing 10 μL of medium. Afterwards, each funiculus was transferred to a 1.5 mL tube containing 40 μL of RNAlater™ Stabilization Solution (Ambion).

### RNA-sequencing library preparation

mRNA was extracted from two biological replicates of 100 funiculi from st 17 of both WT and *stk*, using the Dynabeads^TM^ mRNA DIRECT^TM^ Micro Kit (Invitrogen), according to the manufacturer’s instructions. Sequencing libraries were prepared following the manufacturer’s instructions in the Low Sample protocol, using the TruSeq^®^ RNA Sample Preparation v2 (Illumina Inc.) and sequenced with a NextSeq 500 sequencer (Illumina). Sequencing was performed as reported in (Kadokura *et al*., 2018).

### RNA-sequencing data analysis

Raw reads were cleaned and trimmed using Trimmomatic v.0.38 (Bolger *et al*., 2014). The resulting reads were aligned against the Arabidopsis genome (Araport11) using STAR v.2.7.0 software with default parameters (Dobin *et al*., 2013). Afterwards, featureCounts v.1.6.3 (Liao *et al*., 2014) was used to generate the count matrix and to calculate gene expression values as raw read counts. Count read values were analysed using the DESeq2 package v.1.38.3 (Love *et al*., 2014) from R software v1.30.1. Regularized logarithm (rlog) transformation of the count matrix was used for both principal component plot (PCA) analysis and heatmap clustering. PCA was generated using the *plotPCA* of DESeq2 package and data were visualised with ggplot2 package v.3.4.1 (Wickham, 2016) in R. Heatmap showing clusters of samples was obtained with pheatmap package v.1.0.12 in R and using the top 1000 genes selected based on the mean of the rlog normalised counts. The total number of genes expressed in *stk*, WT or both samples was plotted using a Venn diagram, and *plotMA* function together with ggplot2 were used to show the log_2_FoldChange (log_2_FC) of all genes. To identify differentially expressed genes (DEGs), the raw count data were analysed using the *DESeq* function. A cut-off of p-value, p-adjust value and log_2_FC was applied to select differentially expressed genes: p-value and p-adjust value < 0.05 and a-1 < log_2_FC > 1.

### Gene set enrichment analysis

To determine enriched gene ontology (GO) terms in the *stk* funiculus transcriptome compared to the WT, genes were ranked as: -Log_10_(p-value) multiplied by the sign of the Log_2_FC (Reimand *et al*., 2019). A Parametric Analysis of Gene Set Enrichment (Kim and Volsky, 2005) with Hochberg multi-test adjustment method was performed using the ranked gene list on agriGO v.2.0 [(Tian *et al*., 2017); http://systemsbiology.cau.edu.cn/agriGOv2/, accessed in July 2020], from which biological process and molecular function GO terms with false discovery rate (FDR) < 0.05 were obtained and analysed.

### Enriched networks among DEGs

To uncover functional links between genes, a list containing up-and down-regulated DEGs was upload to GeneMANIA Cytoscape plugin v3.10.0 (Shannon *et al*., 2003) software with a network weighting based on GO biological processes, an FDR < 0.05 and remain settings as default.

### Sample collection, RNA isolation and cDNA synthesis for validation by qPCR

A total of six samples for RNA extraction were collected from funiculi of st 17 flowers, representing two genotypes (WT and *stk*), from three biological samples. One hundred funiculi from one plant were used for each biological replicate. Samples were immediately frozen in liquid nitrogen and stored at -80 °C for follow-up experiments. Total RNA was extracted using the RNeasy Plant Mini Kit (QIAGEN) according to the manufacturer’s protocol, with a minor change: the recommended quantities for reagents were reduced by half. RNA concentration was measured using a spectrophotometer (DS-11 Series Spectrophotometer/Fluorometer). Following the manufacturer’s instructions, 21 ng of total RNA from funiculi of st 17 flowers was used for cDNA synthesis using the Maxima H Minus First Strand cDNA Synthesis Kit (Thermo Fisher). The cDNA products were diluted to 0.21 ng/μL with nuclease-free water prior to qPCR.

### Primer design and qPCR analysis for validation of RNA-sequencing

Primers were designed using Primer3 v.4.1.0 (Koressaar and Remm, 2007; Untergasser *et al*., 2012; Kõressaar *et al*., 2018), following the parameters of (Ferreira *et al*., 2023). For *MLP28*, Litholdo *et al*. (2016) primers were used, and for *ACTIN 7* (*ACT7*), *YELLOW-LEAF-SPECIFIC GENE 8* (*YLS8*) and *HISTONE 3.3* (*HIS3.3*), Ferreira *et al*. (2023) primers were used. Supplementary Table S1 presents all the primers used for qPCR. Primer specificity was confirmed by conventional PCR and electrophoresis on 1% (w/v) agarose gel. qPCR was performed according to Ferreira *et al*. (2023). Briefly, reactions were prepared in a 10 µL final volume containing 5 µL of 2x SsoAdvanced™ Universal SYBR^®^ Green Supermix (Bio-Rad), 0.125 µL of each specific primer pair at 250 nM, 0.75 µL of nuclease-free water, and 4 µL of diluted cDNA template. The reactions were performed in 96-well plates and run on a CFX96 Real-Time System (Bio-Rad) under the following cycling conditions: initial denaturation at 95 °C for 30 s, followed by 40 cycles at 95 °C for 15 s and 60 °C for 30 s, and an additional data acquisition step of 15 s at the optimal acquisition temperature. All reactions were run in three technical replicates and all assays included non-template controls (NTCs). Using three points of a 5-fold dilution series (1:5, 1:25 and 1:125) from a pool of funiculi cDNA (containing both WT and *stk* cDNA), a standard curve was generated to estimate the PCR efficiency of each primer pair using CFX Maestro software v.2.0 (Bio-Rad). The slope and coefficient of determination (R^2^) were obtained from the linear regression line and the amplification efficiency (E) was calculated according to the formula E = (10^−1/slope^ − 1) x 100%. The R^2^ value should be higher than 0.980 and the E value should be between 90% and 110%. After completion of the amplification reaction, melt curves were generated by increasing the temperature from 65 °C to 95 °C, with fluorescence readings acquired at 0.5 °C increments. From the melt curve, the optimal temperature for data acquisition (3 °C below the melting temperature of the specific PCR product) was determined and the specificity of the primers was confirmed (Supplementary Table S2). The sample maximisation method was chosen as the run layout strategy, in which all samples for each defined set were analysed in the same run, thus, different genes were analysed in distinct runs (Hellemans *et al*., 2007). The quantitative cycle, baseline correction and threshold setting were automatically calculated using CFX Maestro software v.2.0 (Bio-Rad). The qPCR products were verified using 1% (w/v) agarose gels. The expression levels of the target genes were quantified in all samples using three reference genes [*ACT7*, *YLS8* and *HIS3.3*; (Ferreira *et al*., 2023)], whose expression stability between *stk* and WT funiculi of st 17 flowers was verified using the RNA-sequencing (RNA-seq) data (Supplementary Table S3). Relative gene expression was calculated using the 2^-ΔΔCt^ method (Livak and Schmittgen, 2001). Data were statistically analysed using the CFX Maestro software v.2.0 (Bio-Rad), which compared the relative expression differences using a Student’s t-test. Statistical significance was set to a p-value threshold of 0.05.

### Construct generation and plant transformation

Genomic regions corresponding to the promoters of three genes (*ADPG1*, *MLP28* and *SCPL41*) were amplified using DreamTaq DNA polymerase (Thermo Fisher) for *ADPG1* and *SCPL41*, and Phusion DNA polymerase (Thermo Fisher) for *MLP28*, using the primer pairs described in Supplementary Table S1. The PCR products were cloned into pENTR™/D-TOPO (Invitrogen) and subcloned into the binary vector pBGWFS7 containing the β*-glucuronidase* (*GUS*) gene (Karimi *et al*., 2002). All constructs were confirmed by DNA sequencing. Expression vectors were delivered into *Agrobacterium tumefaciens* GV3101 (pMP90RK) and were used to transform *A. thaliana* (WT and *stk* genotype) by the floral dip method (Clough and Bent, 1998). Transformant lines were obtained using glufosinate ammonium (BASTA®) as a selection agent.

### Detection of GUS activity

For histochemical analysis, siliques of st 17 flowers from T3 generation plants were fixed in acetone 90% (v/v) (Thermo Fisher) at -20 °C overnight (ON). Samples were then washed twice with sodium buffer [0.2 M sodium dihydrogen phosphate (NaH_2_PO_4;_ Panreac), 0.2 M sodium hydrogen phosphate (Na_2_HPO_4_; Sigma Aldrich)] for 10 minutes and incubated in a X-Gluc solution [50 mM Na_2_HPO_4_, 50 mM NaH_2_PO_4_, 0.2% (v/v) Triton X-100 (Sigma Aldrich), 2 mM potassium hexacyanoferrate(II) trihydrate (C_6_FeK_3_N_6_^+2^ · 3H_2_O; Sigma Aldrich), 2 mM potassium hexacyanoferrate(III) (C_6_FeK_3_N_6_^+3^; Sigma Aldrich), 1 mg/mL X-Gluc (5-bromo-4-chloro-3-indolyl β-D-glucuronic cyclohexylammonium salt; Biosynth)] ON at 37°C. After chemical GUS detection, samples were rinsed with a 90% (v/v) ethanol, followed by 70% (v/v) ethanol for 10 minutes and incubated in a clearing solution [160 g of chloral hydrate (Sigma Aldrich), 100 mL of deionised water, and 50 mL of glycerol (Sigma Aldrich)], where they were kept at 4 °C ON. The following day, siliques were dissected under a stereomicroscope (model C-DSD230; Nikon) using hypodermic needles (0.4 × 20 mm; Braun). The funiculi attached to the septum were maintained in a drop of clearing solution and covered with a cover slip. Samples were observed under an upright Axio Imager AZ microscope (Zeiss) equipped with differential interference contrast optics. Images were captured with an Axiocam MRc3 camera (Zeiss) and processed using Zen 2011 Software (blue edition; Zeiss).

### Image processing

All images were processed for publication using Fiji (Schindelin *et al*., 2012) and Adobe Photoshop CC 2019.

## Results

### RNA-sequencing analysis of *stk* funiculi from st 17 flowers

A procedure for funiculi collection from dissected siliques was developed to address the question of which genes were differentially expressed in the funiculus in the absence of STK. Siliques at st 17 from both *stk* and WT (according to Smyth *et al*.,1990) were harvested, the valves and seeds were removed, and the transmitting tract with the attached funiculi was dipped into an *in vitro* ovule culture medium (Gooh *et al*., 2015) (Fig. 1A-C). Afterwards, each funiculus was cut near the septum and transferred to a clean ovule culture medium (Fig. 1D-E) before being moved to the final tube containing a solution of RNAlater. Each biological replicate was a pool of 100 funiculi from the same plant to ensure that sufficient mRNA was extracted to proceed for Illumina sequencing. The total number of reads as well as the uniquely mapped reads to the *A. thaliana* genome are reported in Supplementary Table S4.

**Fig. 1.**
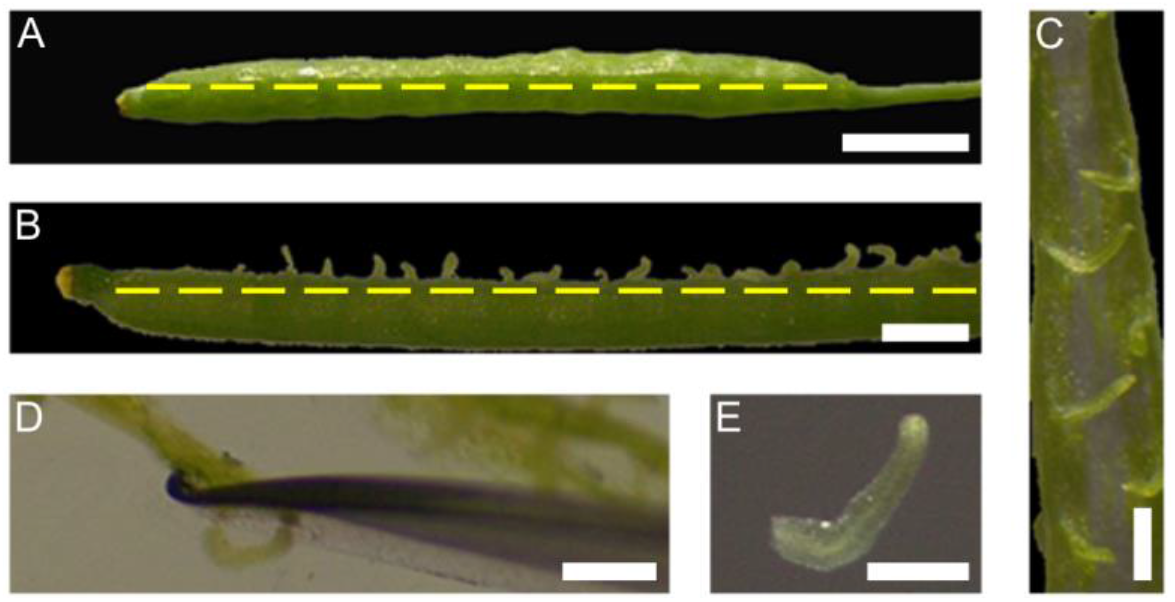
Collection method of funiculi from st 17 flowers for RNA-seq. A) Silique was displayed with one of the valves facing up and a longitudinal cut (yellow dashed line) was performed to remove the valve and the seeds from inside. B) Silique was then placed with the septum facing up and a longitudinal cut (yellow dashed line) was performed along it to remove the other valve and the seeds. C and D) The transmitting tract with the attached funiculi was immersed in an *in vitro* ovule culture medium and each funiculus was cut near the septum with a razor. E) Each individual funiculus was transferred to a new drop of *in vitro* ovule culture medium before being placed inside the collection tube. Scale bar = 1 mm (A and B); 200 μm (C - E).

To investigate if the transcriptional profiles of *stk* and WT funiculi were similar, a principal component analysis (PCA) and hierarchical clustering were conducted (Fig. 2A). PCA showed that 90% of the total variance could be explained by the first two components (53% by PC1 and 37% by PC2). A cluster of *stk* replicates could be identified, but WT replicates showed a slight discrepancy between each other, with one of the replicates being closer to the *stk* replicates. However, the expression analysis of the top 1000 genes clearly showed two distinct clusters, one for *stk* replicates and the other for WT replicates (Fig. 2B). The heatmap also demonstrated that the expression of these genes between *stk* and WT samples was similar, suggesting a low number of differentially expressed genes (DEGs) between the two genotype samples. Overall, the results showed clusters between replicates, even though the expression profiles of *stk* and WT samples were not completely distinguishable.

**Fig. 2.**
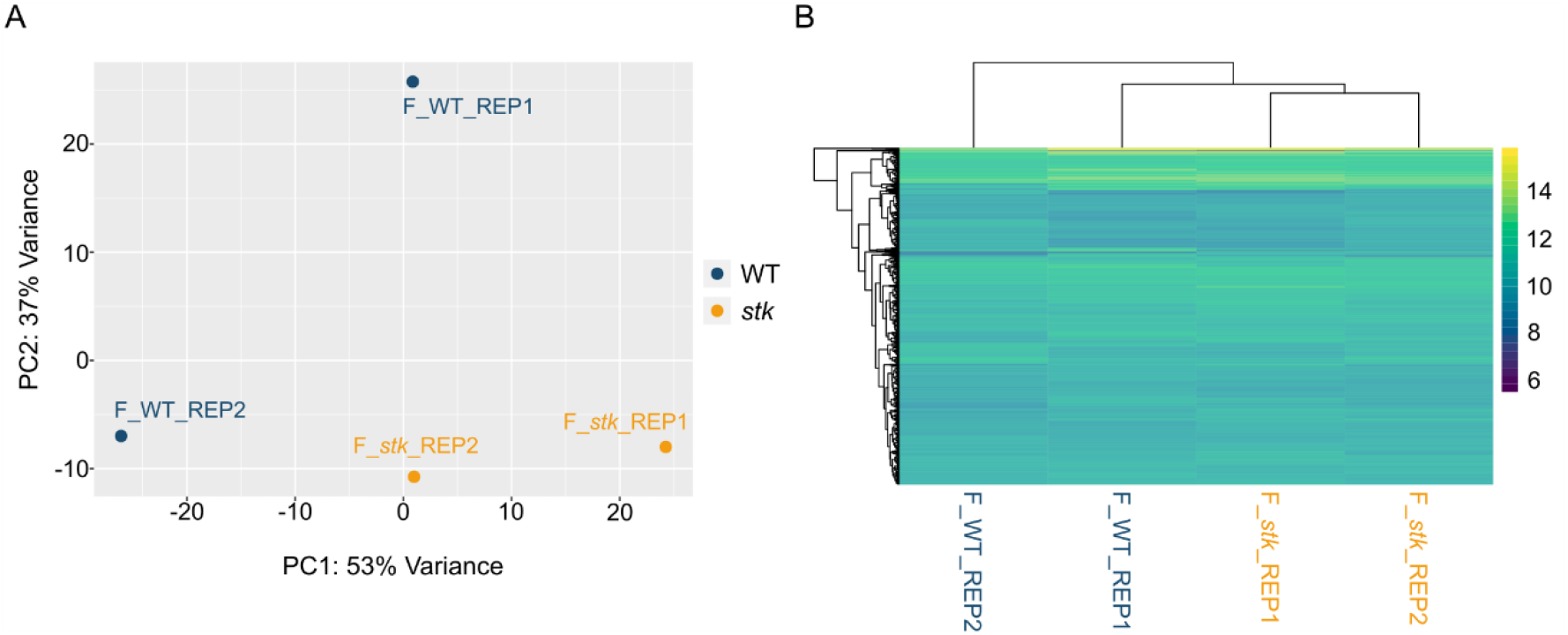
Assessment of samples quality. A) Principal component analysis (PCA) conducted on rlog normalised gene expression values of *stk* and WT samples. X− and Y−axes show PC1 and PC2, respectively, with the amount of variance contained in each component being 53% and 37%, respectively. Each point in the plot represents a biological replicate of 100 funiculi, with a total of four biological replicates in the plot. Symbols of the same colours are replicates of the same experimental group, where orange represents *stk* funiculi, and blue represents WT funiculi. B) Heat map of the top 1000 genes selected based on the mean of the rlog normalised counts. Individual samples are shown in columns and genes in rows. The upper axis shows the clusters of samples, and the left vertical axis shows the clusters of genes. The colour scale represents the relative expression of genes: purple indicates low relative expression levels and yellow indicates high relative expression levels.

To obtain an overview of the funiculi transcriptome, the total number of expressed genes in *stk* and WT samples was examined. Both datasets presented a similar number of genes (21841 for *stk* and 21723 for WT), from which 1293 genes (5.9%) were exclusively expressed in *stk* funiculi, and 1175 genes (5.4%) were only presSent in WT funiculi. The commonly expressed genes expressed in both datasets numbered 20548 (Fig. 3A). Differential expression analysis revealed a total of 169 DEGs (Supplementary Table S5), with p-value and p-adjust value < 0.05 and a -1 < log_2_FoldChange (log_2_FC) > 1, from which 119 genes were upregulated and 50 were downregulated, as seen in the MA plot (Fig. 3B).

**Fig. 3.**
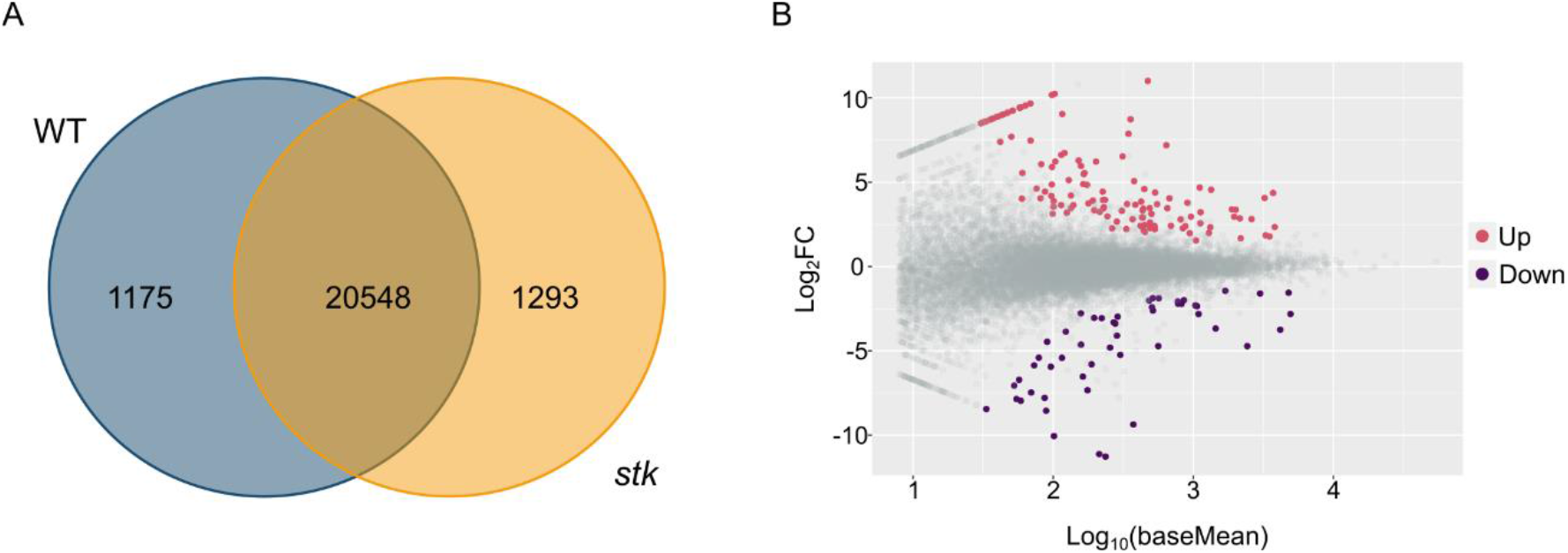
Global analysis of *stk* vs. WT transcriptome from funiculi of st 17 flowers. A) Venn diagram showing the number of genes expressed in each genotype. The number of genes expressed only on *stk* funiculi or WT funiculi are shown, as well as the number of overlapping genes between transcriptomes. WT is represented by a blue circle and *stk* by an orange circle. B) MA plot demonstrates the relationship between the log_10_ average normalised expression on the x−axis and the significance of the differential expression test expressed as log_2_FoldChange (log_2_FC) on the y−axis for each gene in the transcriptome. It illustrates the number of DEGs in the *stk* funiculi. Gray dots represent genes that are not significantly differentially expressed, while pink and purple dots represent genes that are significantly up-and downregulated, respectively, with p-adjust value < 0.05.

### Parametric Analysis of Gene Set Enrichment

To better understand the processes being affected in *stk* funiculus compared to WT, a parametric analysis of gene set enrichment (PAGE) was performed. PAGE is a statistically sensitive method that considers gene expression levels to rank an annotated gene cluster. Genes were ranked following the formula: -Log10(p-value) multiplied by the sign of the Log_2_FC. Each gene ontology (GO) term has an associated z-score value, which provides information about the significance of the enriched term among the data. Moreover, the z-score can assume a positive or negative value, depending on whether the term includes more up-or downregulated genes, respectively. For a full list of biological and molecular function GO terms, please see Supplementary Table S6. Significant enrichment of 53 and seven GO terms for biological function and molecular function, respectively, was observed among the gene datasets. The most significantly upregulated enriched GO category was cellular cell wall organization or biogenesis (GO:0070882) (Fig. 4A), with more than 100 genes associated. Terms related to dehiscence were active in *stk* transcriptome: dehiscence (GO:0009900), lignin biosynthetic process (GO:0009809) and phenylpropanoid metabolic process (GO:0009698). In addition, many cell wall related terms were found such as hemicellulose metabolic process (GO:0010410), xylan biosynthetic process (GO:0045491) and glucuronoxylan biosynthetic process (GO:0010417). Apart from these, regulation of DNA binding (GO:0051101) was a term that stood out. Concerning the molecular function, only two terms were upregulated, oxidoreductase (GO:0016722) and acyl-CoA hydrolase activity (GO:0047617), with the first one presenting a higher fold enrichment (Fig. 4B). PAGE analysis identified the GO term glycoside catabolic process (GO:0016139) as the most inhibited (Fig. 5A). The absence of *STK* in the funiculus affected the sucrose metabolism (GO:0005985), starch metabolism (GO:0005982) and the immune effector process (0002252). Furthermore, the term cell cycle (GO:0007049) appeared in the analysis, containing more than 400 genes, as well as other related terms like posttranscriptional regulation of gene expression (GO:0010608) and gene silencing (GO:0016458). The glutathione metabolic process and gynoecium development were also downregulated in the *stk* funiculi transcriptomic profile. The enriched molecular functions for the downregulated genes revealed terms exclusively related to enzymatic activity: transferase activity, sucrose alpha-glucosidase and beta-glucosidase activity (Fig. 5B).

**Fig. 4.**
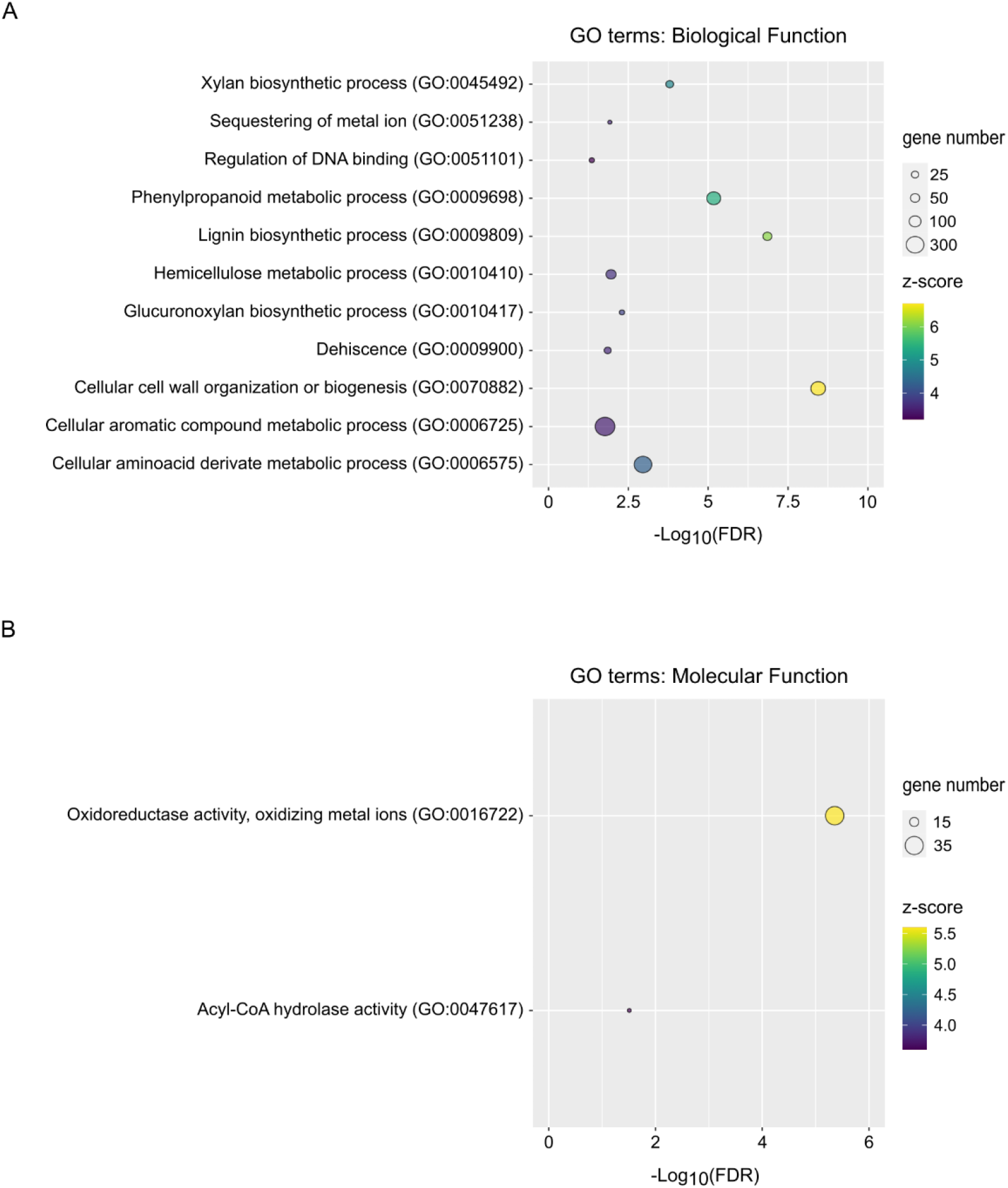
Parametric GO gene set enrichment analysis of activated pathways in *stk* vs. WT transcriptome from funiculi of st 17 flowers. GO enrichment was evaluated at two different levels: A) Biological Processes and B) Molecular Function. Bubble plot diagrams show the significance [-Log_10_(false discovery rate; FDR)] on the x-axis, the fold enrichment (z-score) in a gradient of colour from purple (underrepresented) to yellow (overrepresented), and the number of genes in each pathway.

**Fig. 5.**
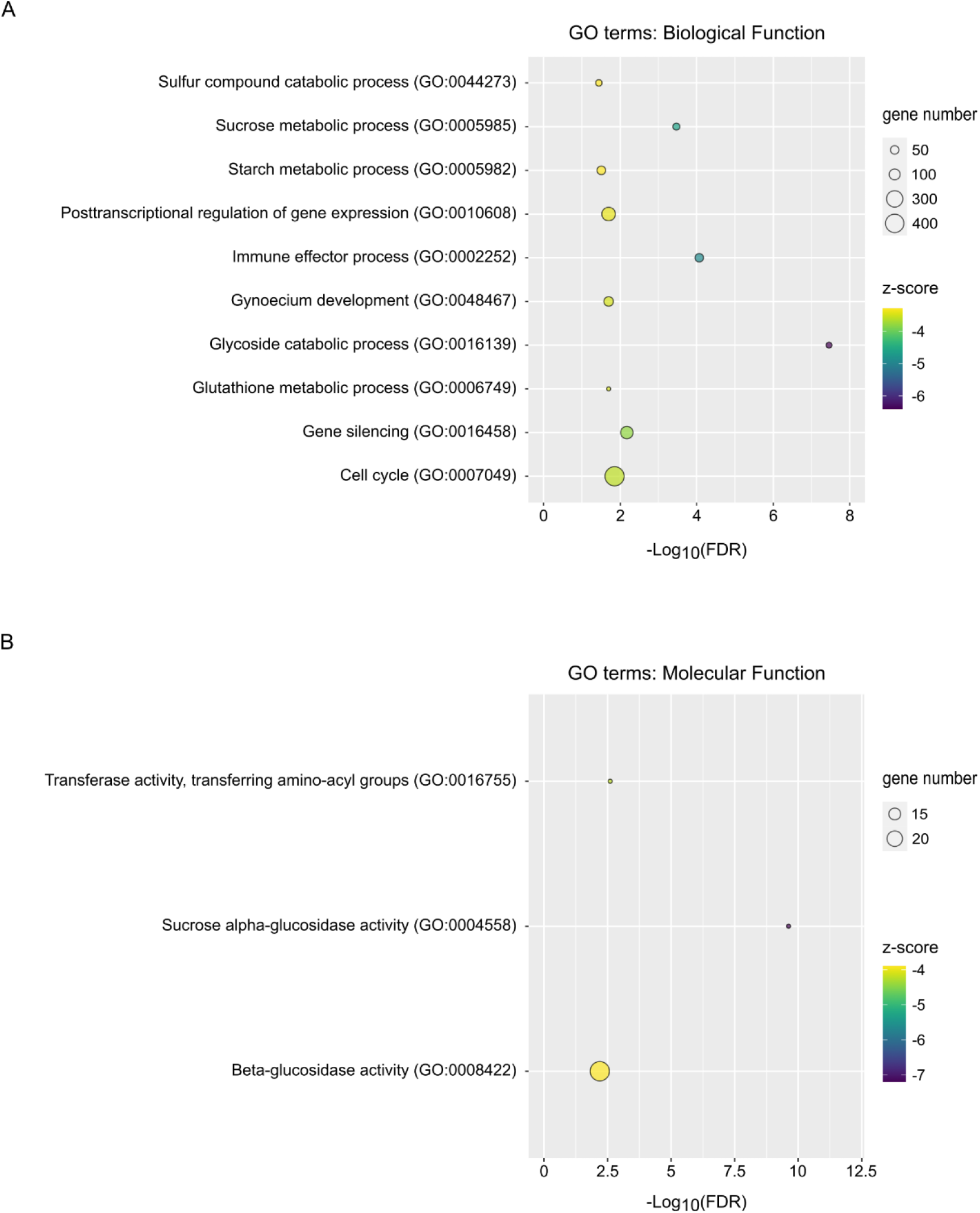
Parametric GO gene set enrichment analysis of inhibited pathways in *stk* vs. WT transcriptome from funiculi of st 17 flowers. GO enrichment was evaluated at two different levels: A) Biological Processes and B) Molecular Function. Bubble plot diagrams show the significance [-Log_10_(false discovery rate; FDR)] on the x-axis, the fold enrichment (z-score) in a gradient of colour from purple (overrepresented) to yellow (underrepresented), and the number of genes in each pathway.

### Expression of funiculus-specific genes in the transcriptome

The expression profiles of genes previously identified as funiculus-specific (Khan *et al*., 2015) were used to assess the reliability of the transcriptome. All 20 funiculus-specific genes were detected in the RNA-seq data (Table 1), with two of them (*EXPANSIN-LIKE B1* and *GRETCHEN HAGEN 3.3*) being downregulated in the *stk* mutant. These genes represent an expansin-like, involved in cell wall extension, and an IAA-amido synthase, implicated in auxin responses, respectively.

**Table 1.** Expression of funiculus-specific genes (Khan *et al*., 2015) on *stk* vs. WT transcriptome from funiculi of st 17 flower. Two genes on this list (*EXPANSIN-LIKE B1* and *GRETCHEN HAGEN 3.3*) were downregulated in the transcriptome (yellow cells). The log_2_FoldChange (log_2_FC), p-value and p-adjust value for each gene can also be seen in the table.

| Locus | Annotation | $\log_2$FC | p-value | p-adjust value |
| --- | --- | --- | --- | --- |
| AT4G17030 | *EXPANSIN-LIKE B1* | -3.75 | 1.05E-09 | 1.22E-06 |
| AT2G23170 | *GRETCHEN HAGEN 3.3* | -1.56 | 1.45E-04 | 2.04E-02 |
| AT2G47160 | *REQUIRES HIGH BORON 1* | -1.92 | 2.53E-03 | 1.71E-01 |
| AT2G22990 | *SINAPOYLGLUCOSE 1* | -1.56 | 2.91E-03 | 1.86E-01 |
| AT2G28305 |  | -1.70 | 1.51E-02 | 5.23E-01 |
| AT2G21220 | Auxin-responsive protein | 1.69 | 2.68E-02 | 7.05E-01 |
| AT5G46240 | *K+ ATPase 1* | -1.65 | 6.09E-02 | 9.53E-01 |
| AT2G22980 | *SERINE*<br>*CARBOXYPEPTIDASE-like*<br>*13* | -2.25 | 6.15E-02 | 9.54E-01 |
| AT5G67420 | *LOB domain protein 37* | 0.71 | 1.69E-01 | 1.00E+00 |
| AT5G37730 |  | -0.32 | 5.91E-01 | 1.00E+00 |
| AT3G43190 | *SUCROSE SYNTHASE 4* | -0.62 | 2.31E-01 | 1.00E+00 |
| AT4G01440 | Nodulin MtN21 family protein | 0.12 | 8.52E-01 | 1.00E+00 |
| AT1G73630 | Calcium-binding protein | 0.30 | 6.95E-01 | 1.00E+00 |
| AT1G68520 | Zinc finger (B-box type) family protein | 0.19 | 8.12E-01 | 1.00E+00 |
| AT1G80760 | *NOD26-like intrinsic protein*<br>*6;1* | 0.52 | 5.50E-01 | 1.00E+00 |
| AT1G64130 | Contains domain Bet v1-like | 0.49 | 5.05E-01 | 1.00E+00 |
| AT1G79430 | *ALTERED PHLOEM*<br>*DEVELOPMENT* | -0.38 | 5.33E-01 | 1.00E+00 |
| AT1G78490 | *CYTOCHROME P450,*<br>*FAMILY 708, SUBFAMILY A,*<br>*POLYPEPTIDE 3* | 0.52 | 5.03E-01 | 1.00E+00 |
| *AT2G15310* | *ADP-RIBOSYLATION*<br>*FACTOR B1A* | -0.99 | 1.96E-01 | 1.00E+00 |
| *AT2G35960* | *NDR1/HIN1-like 12* | -0.40 | 5.74E-01 | 1.00E+00 |

### Enriched networks on DEGs list

To further investigate enriched networks among DEGs and how genes correlated with each other’s regarding physical interactions, co-expression and shared protein domains, the up-and downregulated DEGs were uploaded separately to the GeneMANIA program. All significant biological functions predicted by this tool with false discovery rate (FDR) < 0.05 were listed in Supplementary Table S7. Regarding the downregulated genes in the *stk* transcriptome, the program detected a network related to carbohydrate transmembrane transporter activity (Fig. 6A). *SUGARS WILL EVENTUALLY BE EXPORTED TRANSPORTER 4* (*SWEET4*) and *SWEET7* were the two DEGs identified as involved in this process, being co-expressed and physically interacting with other members of SWEET family. Another network that stood out was associated with floral development (Fig. 6A), containing four DEGs: *CELL WALL INVERTASE 4* (*CWINV4*), *ARABIDOPSIS DEHISCENCE ZONE POLYGALACTURONASE 1* (*ADPG1*), *CYTOCHROME P450, FAMILY 90, SUBFAMILY D, POLYPEPTIDE 1* (*CYP90D1*) and *ARGONAUTE 9* (*AGO9*). However, no co-expression was detected between these genes. These two selected enriched networks share not only a common gene, *SWEET13*, but are also linked by co-expression. For instance, *CWINV4* is co-expressed with *SWEET4*, while *ADPG1* is co-expressed with *SWEET7*. For the upregulated DEGs, the most enriched network with smallest FDR identified by the program was associated with plant type-cell wall biogenesis (Fig. 6B). Nine DEGs were identified such as *NAC DOMAIN CONTAINING PROTEIN 43* (*NAC043*), *NAC012*, *LACCASE 4* (*LAC4*) and *KNOTTED-LIKE HOMEOBOX OF ARABIDOPSIS THALIANA 7* (*KNAT7*). In addition, other DEGs and genes that the program identified as involved with cell wall biogenesis can be visualised. For instance, *WRKY12* is co-expressed with both *NAC043* and *NAC012*, *LAC4* is co-expressed with *LAC2*, which is also a DEG in the present transcriptome, and is co-localised with *KNAT7*.

**Fig. 6.**
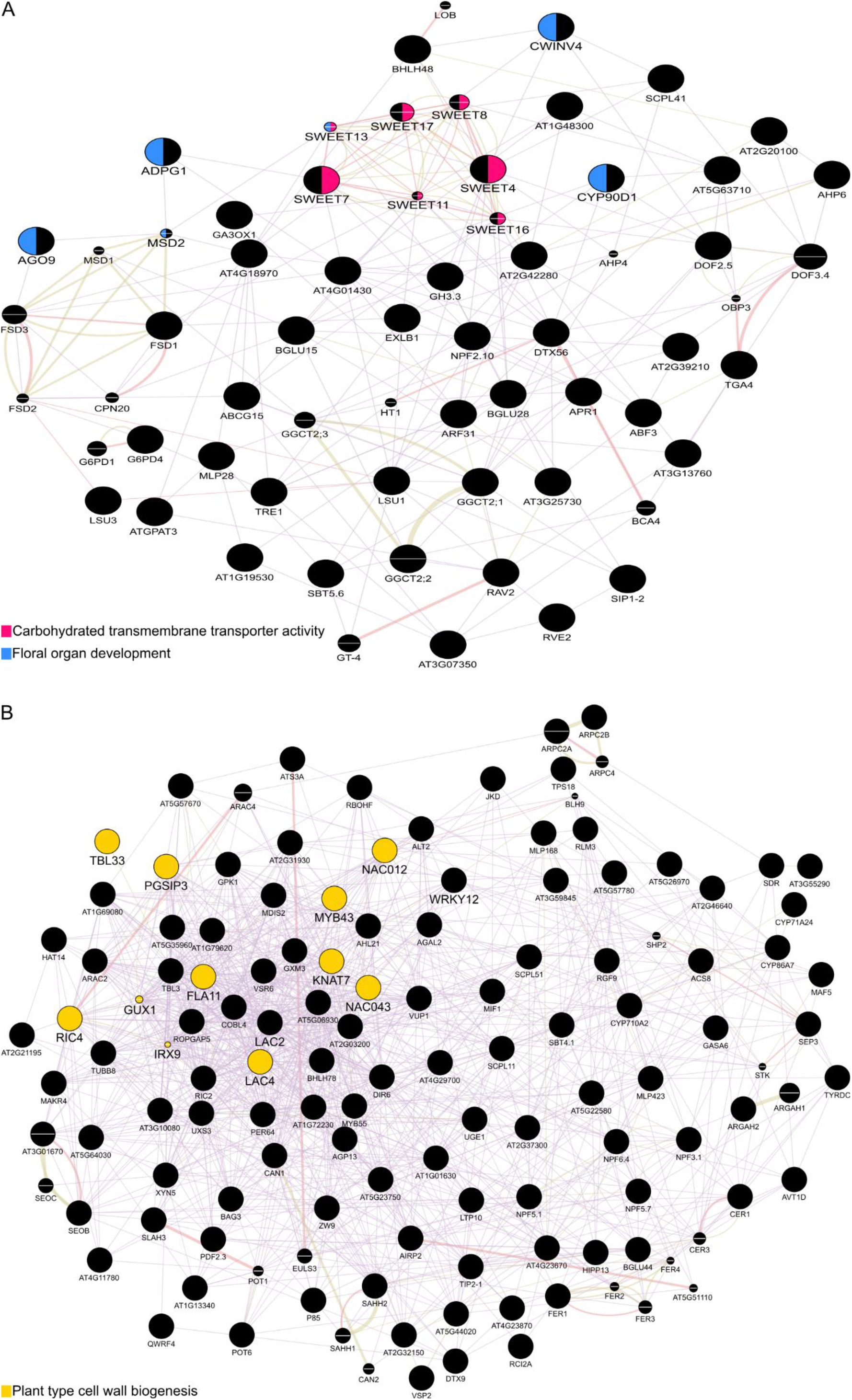
Expression networks of *stk* vs. WT transcriptome from funiculi of st 17 flowers derived from GeneMANIA. A) Networks known from the downregulated differentially expressed genes (DEGs). B) Networks identified from the upregulated DEGs. Genes added as relevant interactors by the GeneMANIA program are shown with a white dash in the middle of the circle, and the size is proportional to the number of interactions they have. The purple edges denote co-expression network interactions, the orange edges represent physical interactions between genes and the green edges indicate shared domain. Different colours in each circle show different functions, as indicated.

### RNA-sequencing data validation through qPCR and promoter expression analysis

To validate the RNA-seq data, we selected a group of up-and downregulated genes from the transcriptome (Table 2), which we considered as potential candidates to be further studied. First, we analysed the GO terms and enriched networks, and we chose genes known to be involved in abscission and seed development. *LAC2* and *LAC4*, members of laccase family, are involved in lignin biosynthesis (Zhao *et al*., 2013; Khandal *et al*., 2020), *SWEET4* and *SWEET7*, belonging to sweet family, are related to sugar transport (Liu *et al*., 2016; Kuanyshev *et al*., 2021) and *ADPG1*, with pectinase activity, is implicated in abscission (Ogawa *et al*., 2009). We also performed a manual analysis on the literature and selected genes from two distinct families. *MAJOR LATEX PROTEIN-like 28* (*MLP28*) and *MLP168* are part of the major latex protein family, which is involved with hormone signalling (Fujita and Inui, 2021), and *SERINE CABOXYPEPTIDASE*-like *41* (*SCPL41*), an enzyme with membrane lipid metabolism functions (-Chen *et al*., 2020). All genes selected were expressed in pistil (Supplementary Fig. S1) and silique (Supplementary Fig. S2) according to the online database ePlant [(Swanson *et al*., 2005); https://bar.utoronto.ca/eplant/, accessed in November 2020) and Arabidopsis eFP Browser [(Mizzotti *et al*., 2018); https://bar.utoronto.ca/efp_arabidopsis/cgi-bin/efpWeb.cgi?dataSource=Silique, accessed in November 2020], respectively.

**Table 2.**
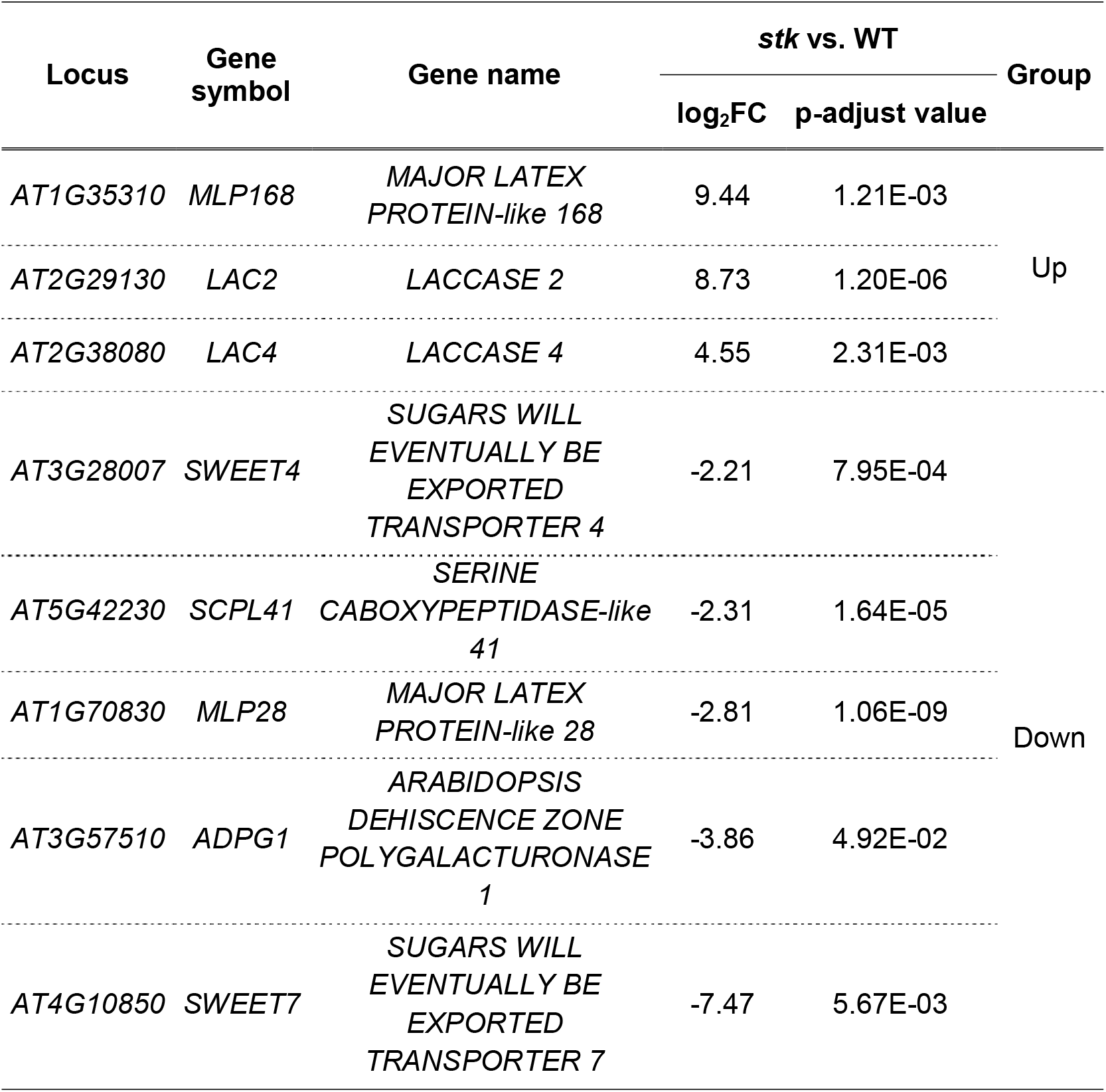
List of differentially expressing genes candidates from the *stk* vs. WT funiculi transcriptome. Gene locus, gene symbol and name, log_2_FoldChange (log_2_FC) and the p-adjust value are detailed for each gene. The list is organised in descending order according to the fold changes.

The relative expression levels of eight selected genes, *MLP168*, *LAC2*, *LAC4*, *SWEET4*, *SCPL41*, *MLP28*, *ADPG1* and *SWEET7* were quantified by qPCR in *stk* and WT funiculi from st 17 flowers (Smyth *et al*., 1990). The transcript levels were normalised to three reference genes (*ACT7, YLS8 and HIS3.3*), previously tested according to Ferreira *et al*. (2023) guidelines to infer about their appropriateness for funiculi samples, and are presented relative to WT funiculi transcript levels (Fig. 7). The analysis revealed that *MLP168*, *LAC2* and *LAC4* expression was higher on *stk* funiculi compared to WT (Fig. 7A). In contrast, *SWEET4*, *SCPL41*, *MLP28*, *ADPG1* and *SWEET7* were downregulated on *stk* samples (Fig. 7B). These results were consonant with the transcriptome dataset (Table 2).

**Fig. 7.**
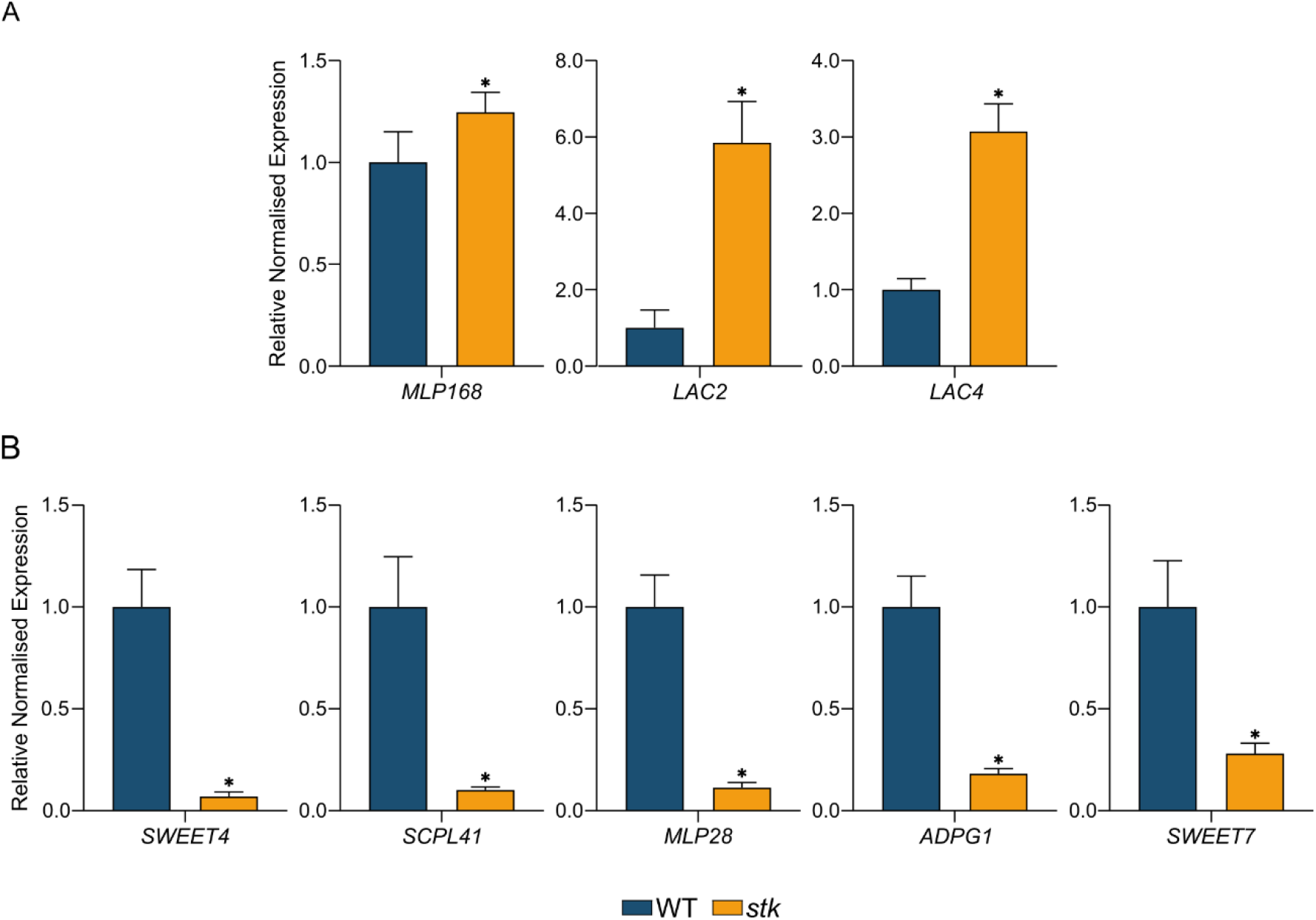
Relative normalised expression levels of *MLP168*, *LAC2*, *LAC4*, *SWEET4*, *SCPL41*, *MLP28*, *ADPG1* and *SWEET7* in WT and *stk* funiculi from st 17 flowers (Smyth *et al*., 1990). A) and B) Relative normalised expression of upregulated and downregulated genes, respectively, in *stk* funiculi according to the RNA-seq data. The transcript levels were normalised to *ACT7, YLS8 and HIS3.3* reference genes (Ferreira *et al*., 2023). The data correspond to the ratio of the expression compared to the WT. Error bars represent the standard error of the mean of three independent biological replicates, each with three technical replicates. Statistical analysis was performed using a Student’s t-test. Asterisks indicate significant differences from WT (*, p < 0.05).

A promoter-driven GUS assay was also performed to further confirm the RNA-seq transcriptome. For instance, we analysed the promoter activity of *MLP28* (p*MLP28*), *SCPL41* (p*SCPL41*) and *ADPG1* (p*ADPG1*). The promoter regions of these genes were fused to the *GUS* reporter gene, and funiculi from both *stk* and WT transgenic plants were observed at st 17 of flower development (Smyth *et al*., 1990), corresponding to the same type of sample used to obtain the transcriptome (Fig. 8). The selection of downregulated genes facilitated the observation of differences in the promoters’ expression between *stk* and WT funiculi. GUS activity driven by p*MLP28* was observed in the entire WT funiculus and septum, whereas in the *stk* background, the detection of the promoter ceased in these two tissues (Fig. 8A and D). In the case of p*SCPL41*, GUS expression was strong in the WT funiculus, however, in the *stk* background, there was no expression of *GUS* in the funiculus (Fig. 8B and E). For p*ADPG1*, the GUS signal was restricted to the seed abscission zone (SAZ) established in the WT funiculus and no GUS staining was visible in the *stk* funiculus (Fig. 8C and F). Overall, these results confirmed the transcriptome data and qPCR, which showed a decrease in transcript levels of *MLP28*, *SCPL41* and *ADPG1* in the *stk* funiculus.

**Fig. 8.**
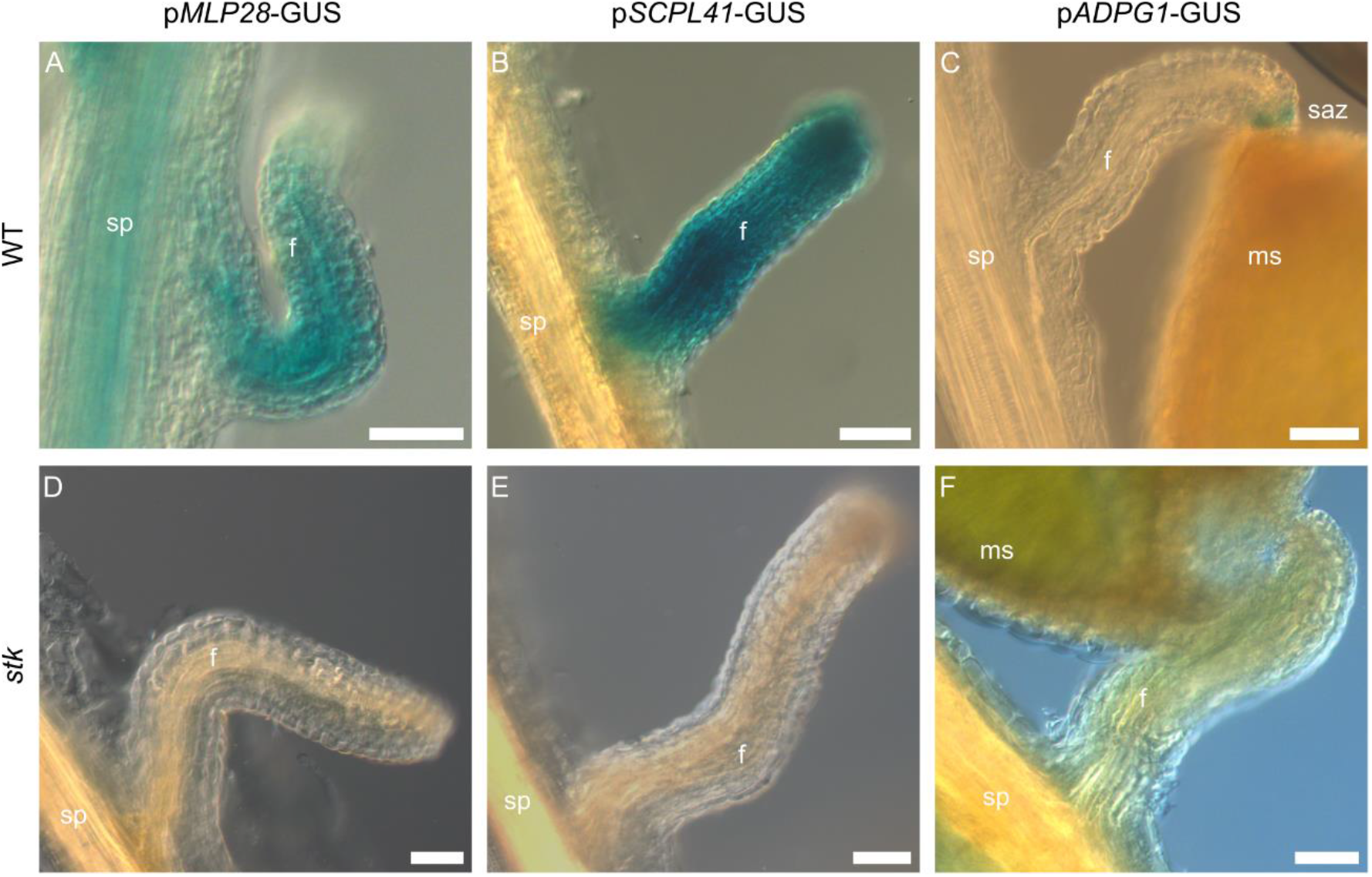
Histochemical localisation of GUS activity in WT (A - C) and *stk* (D - F) funiculi from st 17 flowers (Smyth *et al*., 1990), expressing p*MLP28*-GUS, p*SCPL41*-GUS or p*ADPG1*-GUS constructs. A and D) p*MLP28* expression was detected in WT funiculus but not in the *stk* background. B and E) p*SCPL41* was strongly expressed in WT funiculus, while the GUS activity ceased in *stk* funiculus. C and F) p*ADPG1* signal was restricted to the SAZ of WT funiculus, but it was not detected on the *stk* funiculus. Abbreviations: f – funiculus; ms – mature seed; saz – seed abscission zone; sp - septum. Scale bar = 50 μm.

## Discussion

Seeds are essential for plants, as they harbour the new offspring, and for humans, as a major source of calories. The processes involved in seed development are complex and require an accurate coordination of several regulatory networks established in different reproductive tissues (Becker *et al*., 2014). The funiculus connects the maternal plant to the ovule and lately to the seed, contributing to nutrient transport, mechanical support, and seed positioning. When one of these functions is altered, as observed in *stk* mutants (Pinyopich *et al*., 2003), seed development is disturbed. This influences seed size and viability as well as seed dispersal, ultimately compromising the reproductive success of the plant. Deciphering the genetic networks controlled by STK in the funiculus is, therefore, of major importance in the frame of elucidating how seed development is being disrupted.

Here we analysed the funiculus transcriptional profile of *stk* and WT st 17 of flower development to infer the processes affected by STK absence and gene regulatory networks enriched among the DEGs. We show that cell wall biogenesis, lignification and sugar transport are enriched in the transcriptome and which DEGs are known to be involved in these processes. We then focus on selecting candidate genes for future studies, by validating their expression using qPCR and/or promoter reporter lines.

We started by performing a collection assay to obtain RNA-seq data from *stk* and WT funiculi of st 17 of flower development. A total of 169 genes (p-adjust value < 0.05) were found to be deregulated in the transcriptome (Supplementary Table S5), indicating that the transcriptome profiles of *stk* and WT were not completely distinct, as supported by PCA. Commonly, RNA-seq studies comparing different tissues or developing stages tend to show a more evident separation between different biological samples and, thus, a high number of DEGs (Hofmann *et al*., 2019; Susaki *et al*., 2021; Zhang *et al*., 2023). Moreover, the transcriptomic data generated hereby were from low-input mRNA, which could explain the reduced coverage of low expressed transcripts and, consequently, the detection of low expressed genes as DEGs. Generally, the number of DEGs is slightly lower when compared to libraries obtained from higher amounts of input RNA (Bhargava *et al*., 2014). It is worth noting that all genes classified as funiculus-specific in (Khan *et al*., 2015) were present in the transcriptome, providing a high level of confidence for further analysis.

The most actively enriched function in *stk* funiculi was related to cell wall biogenesis (GO:0070882), consistent with previous phenotypic observations (Pinyopich *et al*., 2003; Balanzà *et al*., 2016). After fertilisation, the WT funiculus vasculature proliferates in a synchronised manner with the cortex and epidermis (Khan *et al*., 2015), with growth being more accentuated in *stk* mutants. Therefore, terms related to the production of cell wall components such as polysaccharides [xylan (GO:0045491), glucuronoxylan (GO:0010417) and hemicellulose (GO:0010410)] were found to be enriched in PAGE analysis. The proliferation of cells is characterised by activation of the transcriptional machinery and, in the case of *stk*, transcription needs to be intensified to allow the overgrowth of the funiculus. This may explain the DNA binding (GO:0051101) GO term enrichment. The *stk* mutant has defects in seed dehiscence (Pinyopich *et al*., 2003) and the reason was later revealed by Balanzà *et al*. (2016), who concluded that an irregular accumulation of lignin in *stk* SAZ caused this phenotype. This control of lignin production by STK was reflected in the enrichment of GO terms for dehiscence (GO:0009900), lignification (GO:0009809) and phenylpropanoid metabolism (GO:0009698), which is a lignin precursor (Yao *et al*., 2021). The analysis of molecular functions revealed the induction of genes related to oxidoreductase activity (GO:0016722). In plants, enzymes such as catalases or peroxidases present this type of activity, and are responsible for regulating the cellular level of hydrogen peroxide, a reduced oxygen species (Pandey *et al*., 2017). In the present transcriptome, *PEROXIDASE 64* (*PRX64*) was upregulated in *stk*, and many studies using loss-of-function mutants or reporter lines observation (Tokunaga *et al*., 2009; Lee *et al*., 2013; Morel *et al*., 2022) have concluded that PRX64 is involved in lignification. The link between this peroxidase and lignification in SAZ has not yet been described, making it an interesting gene to be further analysed. The most inhibitory process in *stk* transcriptome was the glycoside catabolic process (GO:0016139). Glycosides are a wide range group of plant secondary metabolites with multifaceted roles in regulating growth, defense and reproduction in plants (Choi *et al*., 2014; Kytidou *et al*., 2020; Moreira *et al*., 2023). Glycoside hydrolases are a family of enzymes responsible for the removal of specific sugar moieties from glycosides. The mutant analysis of a specific enzyme, BRANCHING ENZYME 1, revealed an impairment in carbohydrate metabolism (Wang *et al*., 2010), which is another GO term inhibited in this transcriptome. Sugars are essential for embryogenesis, and the fact that STK is controlling both glycosylation and sugar catabolism (GO:0005985) may in part explain the small seed phenotype that *stk* present. Also related to *stk* seed size are the terms starch metabolism (GO:0005982) and glutathione metabolism (GO:0006749). Starch is a source of sugar production in the funiculus, that can possibly be transported to the endosperm to nourish the embryo (Khan *et al*., 2015). On the other hand, glutathione is a cell redox that helps in preventing an accumulation of reactive oxygen species within the cell (Gill and Tuteja, 2010). A previous study showed that glutathione is expressed and accumulates in the funiculus as well as it is crucial for proper embryo development and seed maturation (Cairns *et al*., 2006). Collectively, these results suggest that STK controls the metabolism and influx of molecules from the funiculus to the seed, which are important for the correct development of this unit.

A closer look at the DEGs list, using GeneMANIA, allowed the identification of different networks and which specific DEGs were part of them, as well as which genes were being co-expressed, shared protein domains or had a known physical interaction with the DEGs. Not all DEGs were associated with a biological process, which may be a result of an uncharacterised gene or the program itself, since GeneMANIA shows improved outcomes if the input gene list is functionally related. Nonetheless, this analysis contributed for the criteria of selecting genes known to be involved in abscission or seed development. One of the enriched networks for downregulated DEGs was associated with carbohydrate transport activity. It is known that carbon is required for seed growth and the source comes from the maternal plant through the funiculus. Two genes from the transcriptome were identified in this network, *SWEET4* and *SWEET7*, both with glucose transport activity, indicating that STK promotes the sugar efflux required for seed filling (Eom et al., 2015), particularly in the form of glucose instead of sucrose. This finding, combined with other previously reported data (Paolo *et al*., 2021*b*,*a*; Di Marzo *et al*., 2022*b*,*a*), may explain the small seed phenotype that *stk* exhibits (Pinyopich *et al*., 2003). The other network that stood out included terms associated with floral organ development. CWINV4 is an enzyme involved in sugar signalling (Ruan, 2012) that hydrolyses sucrose to generate glucose and fructose. It was first found to be involved in nectar production in Arabidopsis (Ruhlmann *et al*., 2010) and, afterwards, a study reported that CWINV4 together with CWINV2 (mentioned in the paper as CWIN4 and CWIN2) are positive regulators of ovule initiation, through sugar signalling (Liao *et al*., 2020). Remarkably, transcriptomic data obtained from the double mutant transgenic line p*STK*-amiRNA*CWIN24* showed several members of the SWEET family as downregulated DEGs, including *SWEET4* and *SWEET7*, as well as TFs, such as STK (Liao *et al*., 2020). Therefore, it would be valuable to investigate whether STK involvement in sugar efflux is due to the control of CWINV, which would produce sucrose that could be transported to the seed via SWEET transporters. So far, we had confirmed by qPCR that *SWEET4* and *SWEET7* are downregulated in *stk* funiculi from st 17 flowers. Another gene, part of the floral development network, was *ADPG1* that belongs to the polygalacturonase family, which is responsible for disassembling the pectin part of plant cell walls (Caffall and Mohnen, 2009). A genetic approach demonstrated that ADPG1 together with ADPG2 are important for silique dehiscence, whereas the combined action of ADPG1, ADPG2 and QUARTET 2 contributes to anther dehiscence (Ogawa *et al*., 2009). Observations based on transcriptional β-glucuronidase activity were in agreement with the phenotype mentioned above: *ADPG1* was mainly expressed in silique dehiscence zones and in anthers, immediately before anthesis, but also in SAZ. Using the reporter line generated in the present study, we simultaneously confirmed that *ADPG1* promoter is specifically expressed in the SAZ and that GUS activity ceases in the *stk* background (in accordance with RNA-seq and qPCR results). Notably, the authors showed that *ADPG1* expression in the SAZ is disrupted in the *hecate 3* mutant, a TF controlled by STK and involved in SAZ formation (Balanzà *et al*., 2016). Despite the absence of an “obvious defect” in the *adpg1* mutants mentioned by the authors, they did not reveal the type of analysis that was performed. Hence, we believe that these findings point to the involvement of ADPG1 with STK during seed abscission. Among the upregulated DEGs, the most enriched network recognised by GeneMANIA was the plant cell wall biogenesis, in which a laccase family member was identified, *LAC4*. Laccases have been associated with lignin biosynthesis (Wang *et al*., 2015), functioning as activators of lignin production (Berthet *et al*., 2011; Zhao *et al*., 2013). Studies have shown that disruption of LAC4, along with other laccases, leads to a decrease in lignin content of several organs, such as stems, siliques and roots (Berthet *et al*., 2011; Zhao *et al*., 2013). Interestingly, only one study has reported the activity of a specific laccase, LAC2, as a negative regulator of lignin deposition in root vascular tissues under abiotic stress (Khandal *et al*., 2020). LAC2 was identified as being co-expressed with LAC4, and qPCR confirmed that these two enzymes were upregulated in *stk* funiculi. Since *stk* SAZ has an overproduction of lignin (Balanzà *et al*., 2016), these findings suggest that LAC2 and LAC4 participate in the STK-network controlling lignin biosynthesis in the funiculus.

Three additional candidate genes were selected for validation: *MLP168* and *MLP28*, members of the pollen allergen Bet v 1 family (Radauer *et al*., 2008) and the pathogenesis-related protein class 10 as well. Their protein conformation exhibits a hydrophobic cavity capable of binding to ligands, such as cytokinins, brassinolides and secondary metabolites, which MLPs transport to other organs via the plant vasculature, triggering, in some cases, downstream transduction signals in those organs (Fujita and Inui, 2021). Extensive studies on MLPs functions, mostly under biotic and abiotic stresses, have been conducted in several plant species, like *A. thaliana*, *Brassica rapa*, and *Vitis vinifera* (Fujita and Inui, 2021). Regarding MLP168, it was reported an interaction of this protein with the ABA INSENSITIVE 5 protein in a yeast two-hybrid assay (Wang *et al*., 2016). We validated by qPCR that *MLP168* was upregulated in *stk* funiculi. As for MLP28, observations of an amiRNA-*MLP28* line showed defects on vegetative growth with alterations in leaf morphology and shoot apex, dwarfness and eventual premature plant death (Litholdo *et al*., 2016). The present transcriptomic data revealed that *MLP28* was downregulated in *stk* funiculi, which was confirmed by qPCR and promoter-driven GUS activity assays. Due to MLPs nature of binding to plant hormones already implicated in several processes of reproduction (Zuniga-Mayo *et al*., 2023), it seems likely their involvement with ligand transportation from the funiculus to the seed. The last gene selected was *SCPL41*, a serine carboxypeptidase-like protein containing a catalytic triad responsible for cleaving the C-terminal peptide bond in proteins or peptides (Fraser *et al*., 2007). The analysis of *scpl41* mutants has revealed an increase in lipid content in seedlings (-Chen *et al*., 2020). Since *stk* funiculus cells undergo an augmentation in size that must be accompanied by overproduction of cell components, such as lipids, it is reasonable to think that SCPL41 control of lipid content in the funiculus may be regulated by STK. The qPCR results showed a subexpression of *SCPL41* in *stk* funiculi from st 17 flowers and the promoter line showed expression of GUS in the WT funiculus, which was absent in the *stk* background.

Overall, the presented RNA-seq analysis complemented a previous phenotyping analysis of *stk* funiculi by showing GO terms and enriched networks related to cell wall biogenesis, sugar transport and abscission. The results demonstrated that STK is mainly a repressor of cell wall and lignin biosynthesis-related genes, but simultaneously an activator of glucose transporters and seed dehiscence genes acting on the separation layer. The validation of the transcriptome reinforced the reliability of the data obtained and confirmed that genes never associated with funiculus functions are, in fact, deregulated in *stk* funiculus. To our knowledge, this is the first RNA-seq data from *stk* and WT funiculi of st 17 flowers, which makes it a starting point to elucidate, at the molecular level, other protein families involved with STK-network in controlling funiculus functions that can directly affect seed development and dispersal.

## Supplementary data

Supplementary Table S1. Primer sequences.

Supplementary Table S2. Conditions of candidate genes qPCR primers.

Supplementary Table S3. Expression values of reference genes in the *stk* vs. WT funiculi RNA-sequencing.

Supplementary Table S4. Illumina sequencing reads from each *stk* and WT replicate.

Supplementary Table S5. Differentially expressed genes from *stk* vs. WT transcriptome from funiculi of st 17 flowers.

Supplementary Table S6. Gene ontology terms from parametric analysis of gene set enrichment.

Supplementary Table S7. Biological functions of differentially expressed genes predicted by GeneMANIA.

Supplementary Fig. S1. Expression pattern of candidate genes in pistil tissues.

Supplementary Fig. S2. Expression pattern of candidate genes during silique development.

## Author Contributions

MJF, TH, and SC: conceptualisation; MJF, JS, and TS: formal analysis; MJF, JS, HT, TH, and SC: funding acquisition; MJF, JS, and TS: investigation; MJF, and HT: methodology; MJF: project administration; TH, and SC: resources; HT, TH and SC: supervision; MJF, and JS: validation; MJF: visualisation; MJF: writing - original draft; MJF, JS, HT, TH, and SC: writing - review & editing. All authors read and approved the final manuscript.

## Data availability

The raw RNA-seq data generated in this study will be submitted to the Gene Expression Omnibus (GEO) database.

## Acknowledgements

We acknowledge Dr Lucia Colombo for providing *stk* seeds, Dr David Haak and Dr Aureliano Bombarely for assistance with RNA-sequencing analysis, and Ayami Furuta for her technical assistance.

